# Chlamydomonas ATX1 is essential for Cu distribution towards the secretory pathway and maintenance of biomass in conditions demanding cupro-enzyme dependent metabolic pathways

**DOI:** 10.1101/2021.08.18.456897

**Authors:** Keegan L. J. Pham, Stefan Schmollinger, Sabeeha S. Merchant, Daniela Strenkert

## Abstract

Copper (Cu) chaperones, of which yeast ATX1 is a prototype, are small proteins with a Cu(I) binding Mx-CxxC motif, and are responsible for directing intracellular Cu towards specific client protein targets that use Cu as a cofactor. The *Chlamydomonas reinhardtii* ATX1 (CrATX1) was identified because of its high sequence similarity with yeast ATX1. Like the yeast homologue, CrATX1 accumulates in iron-deficient cells (but is not impacted by other metal-deficiencies), and YFP-ATX1 is distributed in the cytoplasm. Reverse genetic analysis using artificial microRNA (amiRNA) to generate lines with reduced CrATX1 abundance and CRISPR/CPF1 to generate *ATX1* knock out lines validated a function for ATX1 in iron-poor cells, most likely because of an impact on metalation of the multicopper oxidase FOX1, which is an important component in high-affinity iron uptake. A more general impact on the secretory pathway is indicated by reduced growth of *ATX1* mutant lines on guanine as a sole nitrogen source, which we attribute to loss of function of UOX1, a urate oxidase involved in guanine assimilation. The block of Cu trafficking towards the secretory pathway in *ATX1* mutants is strikingly evident by a reduced amount of intracellular Cu in all conditions probed in this work.

## Introduction

Copper (Cu) is an important trace element in all kingdoms of life, since it serves as an essential cofactor in enzymes that participate in diverse metabolic pathways, from photosynthesis and respiration to oxidative stress protection and iron acquisition. While a particular quota of Cu is required for fully functioning metabolism, excess intracellular Cu ions may result in damage of macromolecules through redox or oxygen chemistry with negative consequences for the cell. Cu assimilation and sequestration is therefore tightly regulated by orchestrated function of Cu transporters, ligands and chaperones.

A general pathway for Cu metabolism has been developed based on discoveries in several systems. Cu enters the cell through a high affinity Cu(I) uptake system that includes CTR/COPT family proteins (Puig and Thiele, 2002). After entry, cytosolic chaperone proteins (of which ATX1 is a prototype) are responsible for subsequent Cu transfer to key metabolic Cu proteins (Culotta et al., 1997; Rosenzweig and O’Halloran, 2000; Shi et al., 2020). More recently, GSH is implicated as a Cu ion carrier between the importers and metallo-chaperones (Miras et al., 2008).

In *Saccharomyces cerevisiae*, where Cu metabolism is well-dissected, three Cu chaperones compete for Cu entering the cell, delivering them to their specific targets (Valentine and Gralla, 1997). yATX1 is a small protein (73 aa) that delivers Cu to the trans Golgi network for incorporation into membrane-bound and secreted Cu proteins (Lin and Culotta, 1995). yATX1 was discovered originally as a suppressor of oxygen toxicity damage in yeast stains lacking SOD1 function, hence its name Atx for anti-oxidant (Lin and Culotta, 1995). The Cu binding motif, MXCXXC, is present also in many ATP dependent Cu exporters (Cu P-type ATPases). yATX1 transfers Cu from the inner plasma membrane to the secretory pathway where Cu P-type ATPases such as Ccc2 are localized for sequestering Cu into the secretory pathway (Hung et al., 1997; Lin et al., 1997). Ccc2 directly interacts with ATX1, enabling Cu transfer to Ccc2 for translocating Cu coupled to ATP hydrolysis (Arnesano et al., 2002; Pufahl et al., 1997). Notably, all Cu-dependent enzymes from the secretory pathway are loaded with Cu ions in the Golgi. One such enzyme is the cupro-protein FET3, a plasma membrane spanning multicopper ferroxidase, which is required for high affinity iron uptake in yeast (Lin 1997). Accordingly, yeast *ATX1* mutants are unable to load Cu into the active site of FET3 resulting in defective in iron uptake and iron-deficiency (Lin et al., 1997). Yeast *ctr1* mutants defective in Cu import and yeast *ccc2* mutants defective in transferring Cu to the secretory pathway also display an iron-deficiency phenotype that can be rescued by excess Cu (Askwith and Kaplan, 1998; Dancis et al., 1994).

Atx-like Cu chaperones are found with high sequence similarity in most eukaryotes, including mammals, algae and land plants. The human ATX1 homologue (*HAH1*) was shown to rescue the growth defect of yeast *ATX1* mutants in iron poor medium by restoring Cu incorporation into newly synthesized Fet3p (Klomp et al., 1997). HAH1 directly transfer Cu ions to the Cu P-type ATPases ATP7A and ATP7B, which are involved in Cu delivery to the secretory pathway (Banci et al., 2005). Two ATX1 homologues named Cu CHaperone (CCH) and AtATX1 have been analyzed from Arabidopsis. Both proteins, CCH as well as AtATX1, complement the iron conditional growth defect of yeast *axt1* mutants (Himelblau et al., 1998; Puig et al., 2007). Direct interaction with the plant Cu P-type ATPase *RAN1* was established for AtATX1 (Puig et al., 2007). Mutants defective in AtATX1 are hypersensitive to both Cu excess and Cu deficiency conditions (Shin et al., 2012), indicating the importance of this protein for Cu homeostasis.

The genome of the eukaryotic green alga *Chlamydomonas reinhardtii*, referred to as Chlamydomonas hereafter, encodes a single ATX1 homologue, CrATX1. Transcriptional profiling indicates that Cr*ATX1* expression is increased in response to iron deficiency in Chlamydomonas along with other components of Fe assimilation like *FOX1, FTR1, FER1*, and *FEA1*. An orthologous function to yeast ATX1 and human HAH1 was suggested based on functional complementation of the yeast *ATX1* strain and the relationship of multi-copper oxidase CrFOX1 to yFet3 (La Fontaine et al., 2002; Urzica et al., 2012). In this work, we extend the original study to document selective *ATX1* expression and protein accumulation in response to iron nutrition (but not Cu or Zn), localize YFP-tagged ATX1 to the cytoplasm with concentration at the secretory pathway, and assess function in Chlamydomonas by reverse genetics under conditions where cupro-proteins are required: Fe-limitation and guanine as a sole N-source, requiring respectively cupro-enzymes FOX1 and UOX1 encoding a multicopper Fe (II) oxidase and urate oxidase. We conclude that CrATX1 is the cytosolic Cu chaperone responsible for Cu delivery from CTR-like Cu importers and/or GSH at the plasma membrane to the secretory pathway.

## Experimental

### Strains and culture conditions

An miRNA targeting Chlamydomonas ATX1 was designed according to (Molnár et al., 2009; Schmollinger et al., 2010) using the WMD3 tool at http://wmd3.weigelworld.org/. Resulting oligonucleotides ATX-1amiFor: ctagtATGGTCGTCAAGACGGCGAAAtctcgctgatcggcaccatgggggtggtggtgatcagcgctaTTTC-CCCGTCTTGACGACCATg and ATX1amiRev: ctagcATGGTCGTCAAGACGGGGAAAtagcgctgatcac-caccacccccatggtgccgatcagcgagaTTTCGCCGTCTTGACGACCATa (uppercase letters representing miRNA*/miRNA sequences) were annealed by boiling and slowly cooling-down in a thermocycler and ligated into SpeI-digested pMS539, yielding pDS16. pDS16 was linearized by digestion with HinDIII and transformed into Chlamydomonas strain CC4351 by vortexing with glass beads. ATX1 amiRNA strains and *Chlamydomonas reinhardtii* strains CC4533 were grown in Tris-acetate-phosphate (TAP) with constant agitation in an Innova incubator (180 rpm, New Brunswick Scientific, Edison, NJ) at 24°C in continuous light (90 µmol m^-2^ s^-1^), provided by cool white fluorescent bulbs (4100 K) and warm white fluorescent bulbs (3000 K) in the ratio of 2:1, unless stated otherwise. TAP medium with or without iron or nitrogen was used with revised trace elements (Special K) instead of Hutner’
ss trace elements according to (Kropat et al., 2011). ATX1 amiRNA strains were either inoculated from a plate into standard TAP(NH_4_) to repress the artificial micro-RNA or a modified TAP medium, where nitrate was substituted instead of ammonium as the sole nitrogen source, TAP(NO_3_).

### Generation of ctr1 KO strains using LbCpf1/CRISPR

CC425, a cell wall reduced arginine auxotrophic strain, was used for transformation with an RNP complex consisting of a gRNA targeting a PAM sequence in intron1 of *ATX1* and LbCPf1 as described in (Ferenczi et al., 2017) and shown in Figure 5 with the following modifications: Cells were grown to a density of 2 × 10^6^ cells per milliliter and counted using a Coulter counter. Cells (2 × 10^7^) were collected by centrifugation (5 min, 1,500*g*) and respuspended twice in Maxx Efficiency Transformation Reagent (1 mL), followed by resuspension in 230 μL of the same reagent supplemented with sucrose (40 mM). Cells were incubated at 40°C for 20 min. Purified LbCpf1 (80 μM) was preincubated with gRNA (1 nmol, targeting TTTGCGCCGCCCGCAGTGTCCAACG) at 25 °C for 20 min to form RNP complexes. For transfection, 230 μL cell culture (5 × 10^5^ cells) was supplemented with sucrose (40 mM) and mixed with preincubated RNPs and HindIII digested pMS666 containing the *ARG7* gene, which complements the defective *arg7* gene in strain CC425 and thus confers the ability to grow without arginine supplementation in the medium. In order to achieve template DNA-mediated editing, ssODN (4 nmol, sequence containing two stop codons after the PAM target site (GGGGCGGGAGTTGGACACAATCTCAATAGCCTACGTT-GCACCCCTTTGCGCCGCCCGCAGTTAATAGCGTGTCCTGGGAAAGCTGGATGGAGTGGACT-CGTACGAGGTCAGCTTGGAGAA) was added (Figure 5B). The final volume of the transformation reaction was 280 μL. Cells were electroporated in a 4-mm gap cuvette (Biorad) at 600 V, 50 μF, 200 Ω by using Gene Pulser Xcell (Bio-Rad). Immediately after electroporation cells were allowed to recover overnight in darkness without shaking in 5 mL TAP with 40 mM sucrose and 0.4% (w/v) polyethylene glycol 8000 and then plated after collection by centrifugation (5 min at RT and 1650*g*) using the starch embedding method (with 60% corn starch). After 12 d, colonies were transferred to new plates. The following program was used for all colony qPCR reactions: 95°C for 5 min followed by 40 cycles of 95°C for 15 s, and 65°C for 60 s. A first colony PCR using oligos: ATX1screenfor TTGCGCCGCCCGCAGTTAATAG and ATX1seqrev CACTCCTGGCAAAAGCACAG identified candidate strains that might have been successfully edited at the ATX1 locus (which will anneal and hence amplify). Clones that showed successfull qPCR amplification were screened a second time using oligos ATX1seqfor TTGGCGCGTAAGTAATGGTG and ATX1seqrev CACTCCTGGCAAAAGCACAG and amplicons were sequenced using oligo ATX1seqrev, which revealed that we had ssODN mediated gene editing within the second exon of ATX1 in 2 clones that showed introduction of the two in-frame stop codons (Figure 5B).

### Antibody production and protein analyses

Antibodies targeting ATX1 were produced by Covance, Inc. by immunization using the subcutaneous implant procedure for rabbits on a 118-day protocol with synthetic peptide Ac-GVDSYEVSLEKQQAV-VRGKALDPQAC-amide. SDS-polyacrylamide gel electrophoresis (PAGE) of total cell extracts or soluble protein fractions was performed using 20-40 μg of protein for each lane as indicated and transferred in a semi-dry blotter to nitrocellulose membranes (Amersham Protran 0.1 NC). The membrane was blocked for 30 min with 3% dried milk in PBS (137mM NaCl, 2.7mM KCL, 10mM Na_2_HPO_4_, 1.8mM KH_2_PO_4_) containing 0.1% (w/v) Tween 20 and incubated in primary antiserum; this solution was used as the diluent for both primary and secondary antibodies, for 1 h respectively. PBS containing 0.1% (w/v) Tween 20 was used for washing membranes twice for 15 min each time. The secondary antibody, used at 1:6000, was goat anti-rabbit conjugated to alkaline phosphatase. Antibodies directed against ATX1 (1:500) and GFP (1:1000, gift from Rose Goodchild, KU Leuven) were used as indicated.

### YFP Plasmid Construct and Cloning

For YFP tagging, we used the pRMB12 plasmid which was a gift from R. Bock (MPI for Molecular Plant Physiology, Potsdam-Golm), (Barahimipour et al., 2015). ATX1 open reading frames (omitting the start codon ATG) were PCR amplified (Phusion Hotstart II polymerase, ThermoFisher Scientific) from genomic DNA, gel purified (MinElute Gel Extraction Kit, QIAGEN) and cloned in-frame with an N-terminal Venus tag by Gibson assembly into EcoRI-cut pRMB12 to generate pDS35. Primer sequences also introduced a Glycine-Serine (GS) linker between ATX1 open reading frames and YFP and were: ATXfor: cctgggcatggacgagctgatcaagGGTGGTGGCGGTTCTTCTACCGAGGTGGTCCTTA and ATXrev: GTACAGGCGGTCCAGCTGCTGCCAGTTACGAGGACACGAGCTCGGCCTTCTTCCC. pDS35 was linearized using XbaI and transformed into UVM11 (a UV-induced mutant derived from CC4350 (*cw15 arg7-8 mt*+) known to efficiently express nuclear transgenes (Neupert et al., 2009) that was kindly provided by R. Bock (MPI for Molecular Plant Physiology, Potsdam-Golm) using the glass bead method (Kindle et al., 1989). Transformants were selected on 10 µg/µl paromomycin and screened by immuno-detection using anti-GFP.

### Immunofluorescence and live cell imaging

IF was carried out as described (Uniacke et al., 2011). In brief, cells were added to glass coverslips coated with Poly-L-lysine for 3-4 min. Excess cells were washed off with 1xPBS for a total of two times. Coverslips were placed in chilled (−20° C) methanol for 10 min, and air dried for 15-20 min. Fixed cells were rehydrated by adding 1xPBS for 20 min. Cells were blocked for 1h at RT in blocking buffer (1% BSA, 0.1% Tween-20 in PBS). Primary antibodies were used in blocking buffer (GFP, 1:100) and were incubated overnight at 4 °C. After removal of the primary antibody solution, cells were wash three times for 5 min with blocking buffer. Cells were incubated in secondary antibody (Alexa 568, 1:100), diluted in blocking buffer for 1 h at room temperature. After three washes for 5 min with blocking buffer, cover slips were mounted with 60% glycerol, TRIS-HCl, pH 7.4.

For live cell imaging of YFP-ATX1 expressing lines, cells were imaged on a Zeiss LSM 880 using 514 Ex and 527 nm emission. Image processing was performed using the Zeiss ZEN software.

### Quantitative metal, phosphorus and sulfur content analysis

1 × 10^8^ cells (culture density of 3-5 × 10^6^ cells/ml) were collected by centrifugation at 2450*g* for 3 min in a 50 ml Falcon tube. The cells were washed 2 times in 50 ml of 1 mM Na_2_-EDTA pH 8 (to remove cell surface–associated metals) and once in Milli-Q water. The cell pellet was stored at -20°C before being overlaid with 286 µl 70 % nitric acid and digested at room temperature for 24 h and 65 °C for about 2 h before being diluted to a final nitric acid concentration of 2 % (v/v) with Milli-Q water. Complementary aliquots of fresh or spent culture medium were treated with nitric acid and brought to a final concentration of 2 % nitric acid (v/v). Metal, sulfur and phosphorous contents were determined by inductively coupled plasma mass spectrometry on an Agilent 8900 Triple Quadropole instrument by comparison to an environmental calibration standard (Agilent 5183-4688), a sulfur (Inorganic Ventures CGS1) and a phosphorus (Inorganic Ventures CGP1) standard. 89Y served as an internal standard (Inorganic Ventures MSY-100PPM). The levels of analytes were determined in MS/MS mode. ^23^Na, ^24^Mg, ^31^P, ^55^Mn, ^63^Cu and ^66^Zn analytes were measured directly using He in a collision reaction cell. ^39^K, ^40^Ca and ^56^Fe were directly determined using H_2_ as a cell gas. ^31^P and ^32^S were determined via mass-shift from 31 to 47 or 32 to 48 utilizing O_2_ as a cell gas. An average of 4 technical replicate measurements was used for each individual biological sample. The average variation between technical replicate measurements was less than 2 % for all analytes and never exceeded 5 % for an individual sample. Triplicate samples (from independent cultures) were also used to determine the variation between cultures. Averages and standard deviations between these replicates are depicted in figures.

## Results

### ATX1 is conserved in most algae

Since many algae have reduced Cu quotas (Omelchenko et al., 2010; Wolfe-Simon et al., 2005), we sought to understand the role of Cu chaperones in Cu homeostasis pathways in green algae. We recovered ATX1 sequences from various eukaryotic organisms using yATX1 as a query. A homolog was found in most algae, indicating the selective pressure for maintaining this molecule even if some algae can grow in Cu-deficient medium by dispensing with abundant Cu proteins (Blaby-Haas and Merchant, 2017). Generally, the yATX1 homolog occurs as a single copy gene, which is in contrast to land plants where at least two distinct ATX1 homologs are found in addition to the Cu chaperones for Cyt oxidase and superoxide dismutase (Himelblau et al., 1998; Puig et al., 2007). All algal ATX1 proteins share the MXCXXC domain that is required for Cu binding and that is a defining characteristic of the ATX1 family of Cu chaperones (Figure 1).

**Figure 1:**
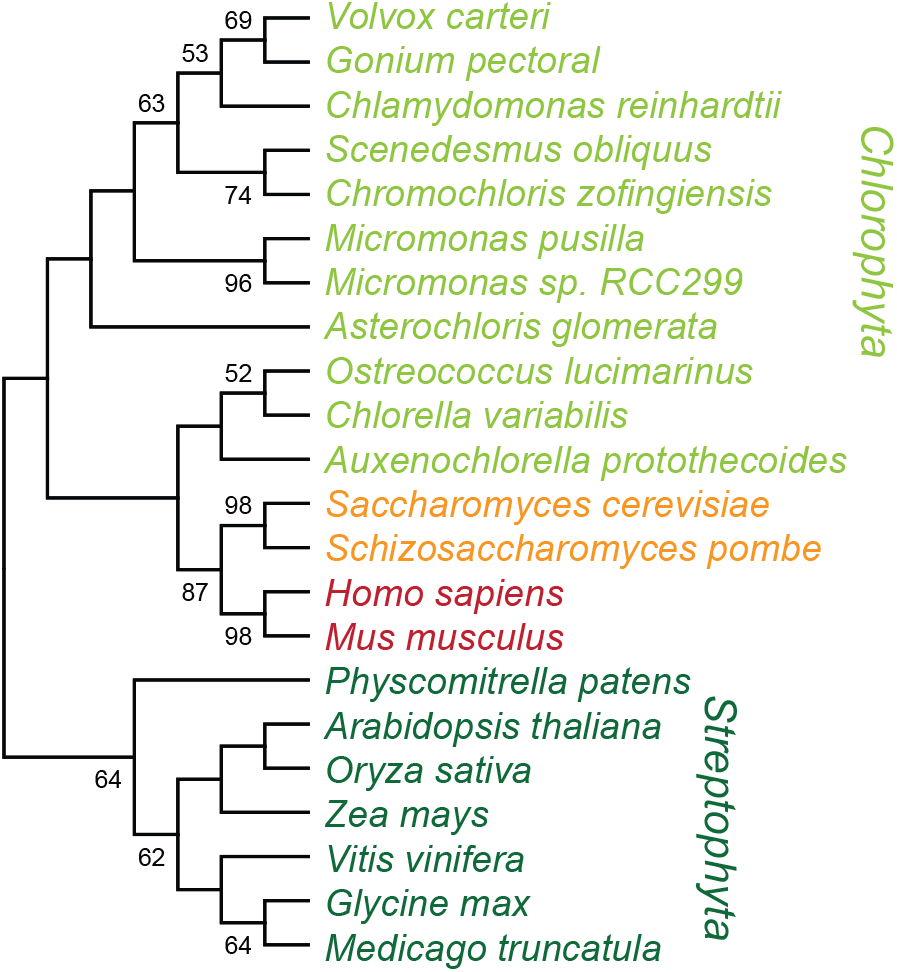
Atx1 is a conserved and widespread protein. The evolutionary history was inferred by using the Maximum Likelihood method and JTT matrix-based model (Jones 1992). The bootstrap consensus tree inferred from 1000 replicates (Felsenstein 1985) is taken to represent the evolutionary history of the taxa analyzed (Felsenstein 1985). When the percentage of replicate trees in which the associated taxa clustered together in the bootstrap test (1000 replicates) exceeded 50% they are shown next to the branches [3]. This analysis involved 22 amino acid sequences. There were a total of 90 positions in the final dataset. Evolutionary analyses were conducted in MEGA X (Kumar 2018).

### ATX1 is predominantly expressed in conditions in which iron is scarce

Based on the sequence relationship of CrATX1 with the yeast and Arabidopsis proteins and the concomitant increases in *ATX1* and *FOX1* transcript abundances under low-iron conditions (La Fontaine et al., 2002), we hypothesized that CrATX1 functioned in Cu delivery to the multi-copper oxidase and hence proposed that ATX1 is involved in Cu(I) routing towards the secretory pathway as in other organisms. A survey of *ATX1* expression in large-scale RNA-seq datasets from experiments involving Fe, Cu and Zn depletion (Castruita et al., 2011; Hong-Hermesdorf et al., 2014; Urzica et al., 2012) confirmed that *ATX1* is highly and selectively responsive to poor iron nutrition (Figure 2A). Indeed, transcripts increase already under asymptomatic Fe-deficiency and are further increased in the Fe-limited symptomatic state. The pattern of expression is unaffected by carbon source (CO_2_ vs. acetate) nor by other trace nutrient deficiencies like Cu or Zn. In general, a survey of 67 RNA-Sequencing datasets indicated that *ATX1* transcript abundances are quite stable across various nutrient regimes and other stress conditions (data not shown), with Fe nutrition being one notable exception. *FOX1* transcripts, on the other hand, are increased not only in Fe deficiency but also in Cu deficient conditions and are reduced in Zn deficient cells (Figure 2A). *ATX1* and *FOX1* are therefore not strictly co-expressed; simultaneous expression is limited to conditions involving reductions of Fe in the growth media.

**Figure 2:**
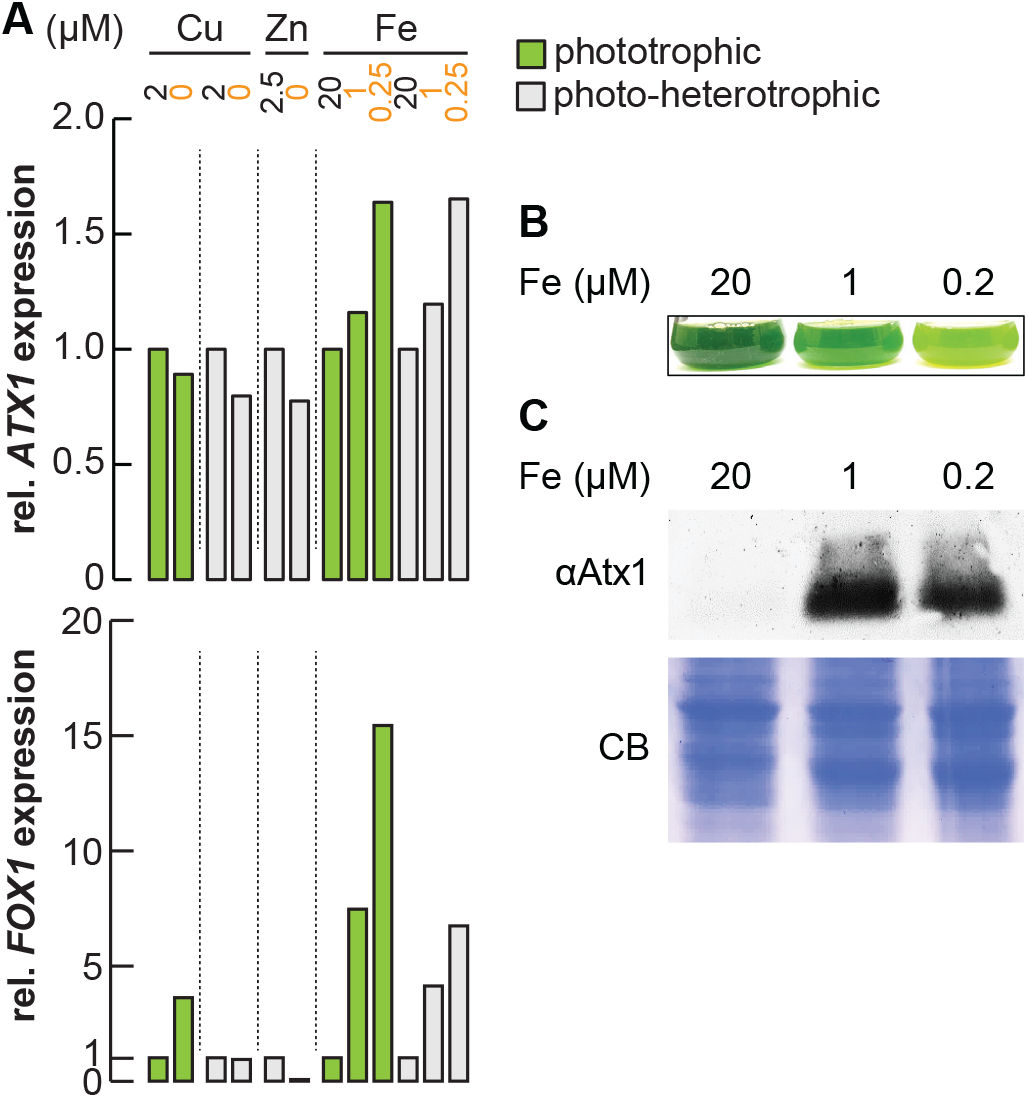
Atx1 expression depends on iron nutritional status. **A:** Relative abundance changes of *ATX1* (upper panel) and *FOX1* (bottom panel) transcripts upon trace metal limitation (Cu, Zn or Fe). Horizontal lines separate different experiments. Green bars indicate experiments under phototropic conditions, grey bars indicate presence of acetate in the light. **B**. *Chlamydomonas reinhardtii* cells CC4533 were grown in either iron replete TAP medium (20), in Fe deficient (1) or in Fe limited (0.2) growth medium as indicated. A picture of flasks was taken 5 days post inoculation. **C**. Total soluble protein was separated by 15 % SDS PAGE and immuno-detected using antisera against Atx1. Coomassie blue (CB) stain was used as loading control. Shown is one of two experiments performed with independent cultures.

To test the impact of the change in mRNA abundance, we analyzed the abundance of the corresponding ATX1 polypeptide as a function of iron nutrition. Chlamydomonas cells were grown in TAP medium supplemented with 20 µM, 1 µM, and 0.2 µM, corresponding to iron -replete, -deficient, and -limited growth respectively (Glaesener et al., 2013). We recapitulated previously-reported phenotypic differences in growth, including chlorosis and growth arrest on media containing 1 and 0.2 µM Fe as compared to 20 µM Fe (Figure 2B). Chlorosis was particularly strong in iron limited cultures, likely due to complete exhaustion of the iron source from the growth medium (Page et al., 2012). Immunoblot analysis indicated that ATX1 is iron-conditionally expressed (Figure 2C), which is consistent with a role for Chlamydomonas ATX1 in Cu delivery to the secretory pathway for FOX1 biosynthesis.

### ATX1 is a cytosolic protein

To distinguish ATX1 localization, we tagged the protein with a fluorescent reporter (YFP) at the N-terminus and expressed the YFP-ATX1 fusion protein from a strong, constitutive promoter (*PSAD*, Figure 3A) into strain UVM11, which expresses nuclear transgenes efficiently (Neupert et al., 2009). We screened soluble cell extracts of 13 transformants by immuno-detection to identify 4 independent lines that showed a signal against both ATX1 and GFP antiserum with a band in each case at the expected size (29 kDa) of the fusion protein (Figure 3B). The analysis using the GFP antiserum showed no other visible bands below the size of the fusion protein, indicating that there were no cleavage products, or expression of YFP without ATX1 in the transformants, which would potentially interfere with localization analysis using confocal microscopy (Figure 3B). We chose YFP-ATX1 line #8 and a line expressing YFP alone (Figure 3) for further analyses, using direct live cell imaging. Notably, using chlorophyll auto fluorescence to visualize the chloroplast, we conclude that the YFP-ATX1 fusion protein is localized to the cytoplasm, including the vicinity of the plasma membrane, consistent with trafficking from the inner plasma membrane to the trans-Golgi network (Figure 4A).

**Figure 3.**
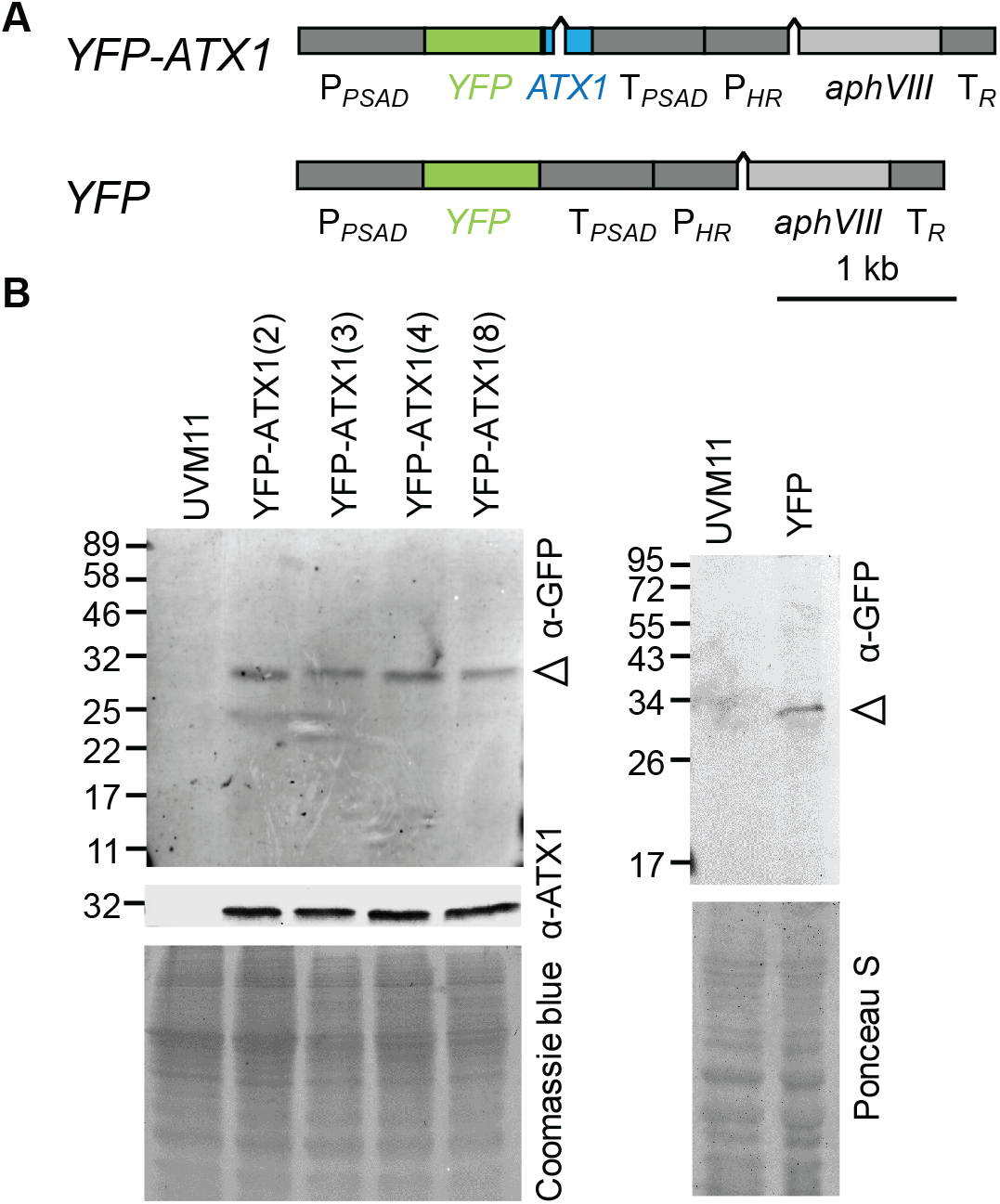
YFP-Atx1 is expressed in Chlamydomonas. **A**. Physical map of the YFP-ATX1 and the YFP control construct. **B**. Total soluble protein from UVM11 (parental strain) and 4 independent UVM11 transformed strains expressing the N-terminal YFP-ATX1 fusion protein as well as YFP were separated by 15% SDS PAGE and after transfer to a nitrocellulose membrane probed using a GFP antibody. ATX1 antibody was also tested in the assay to confirm cross reactivity with the fusion protein. Coomassie blue stain (CB) of the gel is shown as a loading control. A white arrow marks the expected size of the YFP-ATX1 fusion protein.

**Figure 4:**
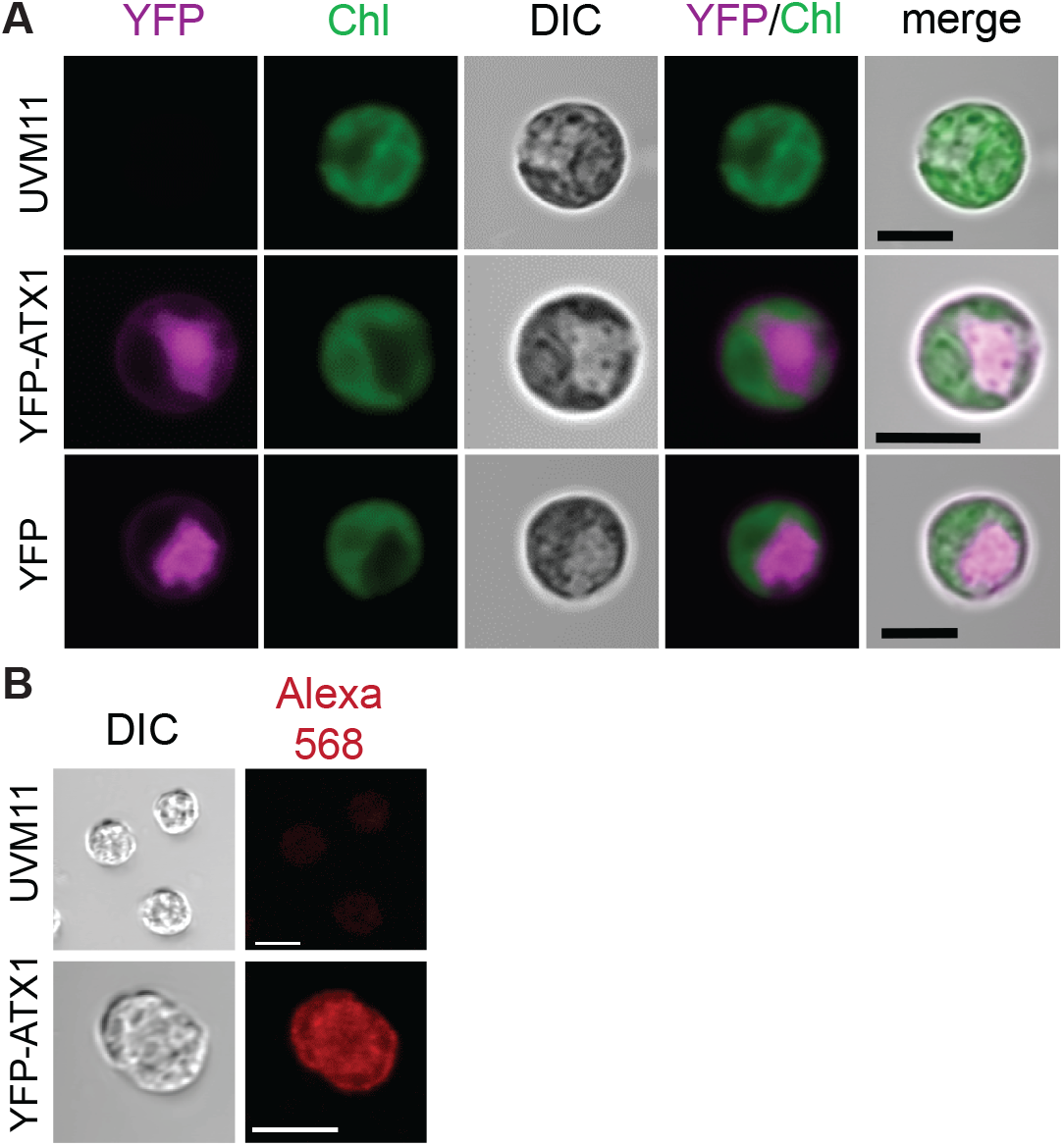
Subcellular localization of YFP-Atx1 fusion protein. **(A)** Confocal microscopy of UVM11 control (untransformed), ATX1 protein fused with Venus (magenta) and Venus expressed without a fusion protein driven by the constitutive *PSAD* promoter, respectively. Chlorophyll auto fluorescence is shown in green (Chl) and bright field images (DIC) are shown to depict the cell and the chloroplast, respectively. All strains were grown in medium supplemented with 1µM Fe. Scale bars correspond to 5 μm. **(B)** Immunofluorescence images of methanol fixed UVM11 control and YFP-ATX1 expressing cells stained with primary antibodies to GFP and secondary antibodies to Alexa568 (red) acquired at equal exposure. Scale bars correspond to 5 µm.

Live cell imaging using YFP fusion proteins is a facile method for assessment of protein localization, with the caveats that YFP signals can be quenched rather rapidly and that live cell imaging is limited to a short period of time due to cell death. Therefore, we recapitulated the result from live cell imaging using fixed cells and indirect immunofluorescence (Uniacke et al., 2011). Indirect immunofluorescence relied on the anti-serum directed against GFP. Cells were sampled on poly-L-lysine-coated coverslips, blocked, and probed with anti-GFP and Alexa Fluor 568 conjugate secondary antibody before fluorescence microscopy. Again, we noted a distinct fluorescence signal inside the cells and at the plasma membrane region in cells expressing the YFP tagged protein, confirming that ATX1 is trafficked to the inner plasma membrane in Chlamydomonas, presumably to accept Cu from GSH and CTRs. We fortuitously identified a pair of dividing cells that clearly show signal at the periphery of the cells as well as around the two forming nuclei, which is evident being in between the two dividing daughter cells (Figure 4B), indicative of TGN localization, again consistent with the findings in yeast.

### ATX1 is required for growth in low iron conditions

To validate the function of ATX1 in Chlamydomonas, we designed an artificial microRNA (amiRNA) targeting the 3’ coding sequence of the Cr*ATX1* gene (Figure 5A, red bar) (Molnár et al., 2009; Schmollinger et al., 2010). We initially screened candidate *ATX1* knock-down lines based on reduction of *ATX1* mRNA as compared to control lines, which were transformed with an empty vector that lacks the target region against CrATX1 (*ATX1*). The candidate knock-down lines were grown in 0.2 µM Fe to maximally induce *ATX1* expression to improve signal to noise in the screen. We chose two lines that showed reduced *ATX1* mRNA abundance, which are named ami-*atx1*(1) and ami-*atx1*(2) (Figure 5B). Compared to empty vector-transformed control strains, the abundance of ATX1 was reduced to about 50% and 25% in Fe-limited ami-*atx1*(1) and ami-*atx1*(2), respectively (Figure 5C). In parallel, we generated also *atx1* knock-out lines using CRISPR/CPF1 mediated gene editing adapted from previous protocols using homology-directed DNA replacement and the enzyme LbCPF1 in combination with a selection marker (Ferenczi et al., 2017; Greiner et al., 2017). *ATX1* has a PAM (<u>P</u>rotospacer <u>A</u>djacent <u>M</u>otif) target sequence for LbCPF1 recruitment within its intron (Figure 5, green bar). We used single stranded oligodeoxynucleotides (ssODNs) as a repair template to introduce two in-frame stop codons within the second exon of the *ATX1* gene (Figure 5B). Sequencing of the PCR product spanning the gene editing site confirmed the presence of the two stop codons in two *ATX1* lines, depicted *atx1-1* and *atx1-2* (Figure 5B).

**Figure 5:**
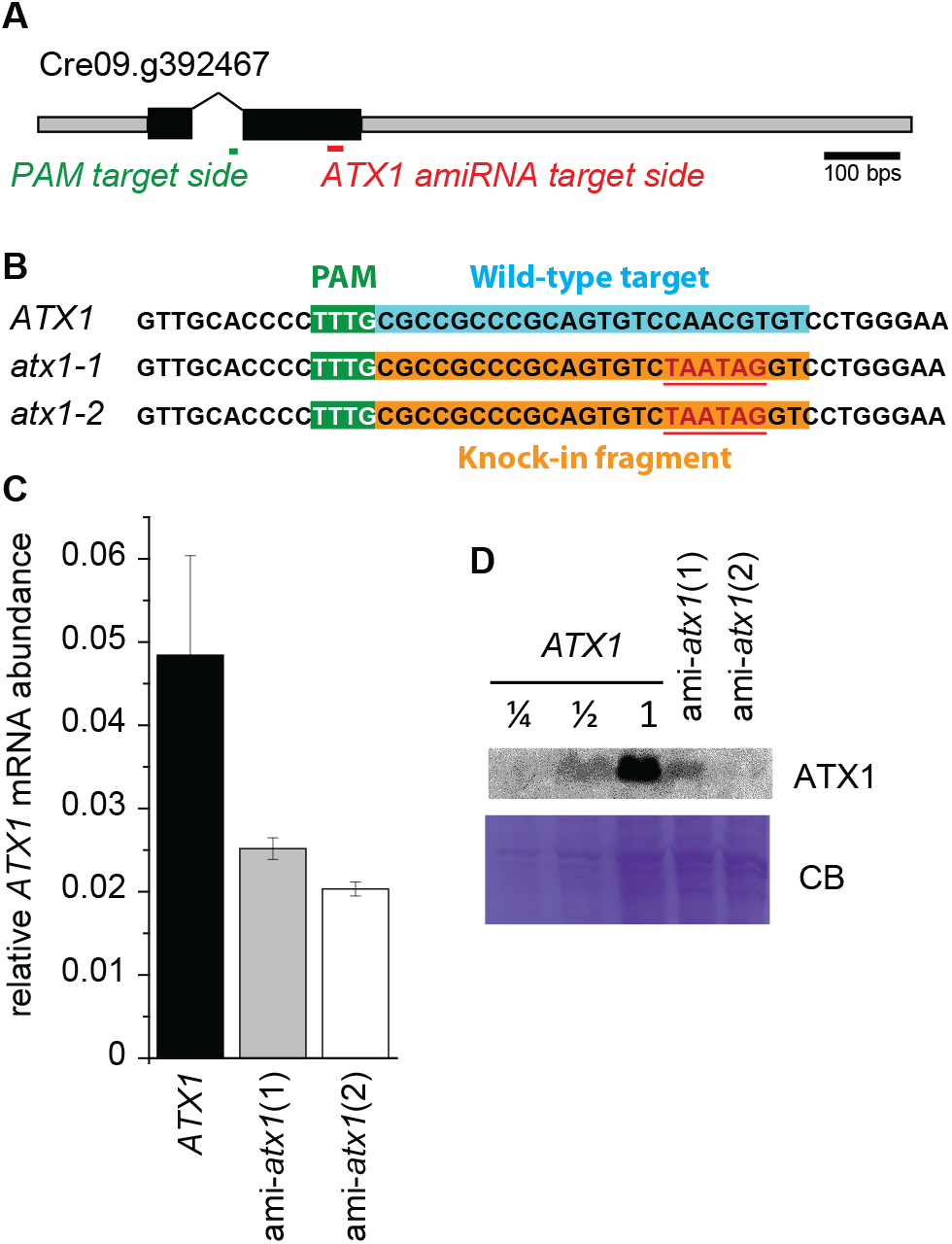
Strains with reduced ATX1 abundance using artificial microRNA constructs. **A**. Physical map of the *ATX1* gene and target site for the amiRNA. **B**. amiRNA lines 1 and 2 grown in TAP supplemented with 0.2 µm Fe (limited) showed reduced *ATX1* mRNA abundance in comparison to Fe limited reference strains that were transformed with an empty vector (*ATX1*). **C**. Soluble protein from Fe limited cultures was separated by 15% SDS-PAGE, transferred to nitrocellulose membranes, and immuno-detection was performed using antisera against ATX1 to confirm reduced ATX1 protein abundance in ATX1 amiRNA lines.

If ATX1 functions in Cu trafficking towards the secretory pathway, *atx1* mutant lines would be affected in FOX1 metalation and hence display a phenotype in iron-poor medium where FOX1 function is required (Terzulli and Kosman, 2009). To test this, we grew *atx1-1*, ami-*atx1*(1), ami-*atx1*(2) and corresponding control lines in iron-replete, -deficient and -limiting media. Neither the amiRNA lines nor *atx1-1* showed a growth defect in iron replete conditions, which is expected since ATX1 as well as its proposed client FOX1, are barely expressed in this situation (Figure 6). On the other hand, in iron-limiting conditions, both ami-*atx1* lines as well as the *atx1-1* mutant showed reduced growth as compared to control lines (Figure 6). When we measured the Fe content of each strain by ICP-MS/MS, we noted that all strains showed a reduction of the Fe content in Fe-poor conditions. Nevertheless, since Fe is a growth-limiting nutrient, we did not see any significant differences in the iron content between reference wild-type, *atx1-1* mutant and ami-*atx1* lines (Figure 7A), as expected.

**Figure 6:**
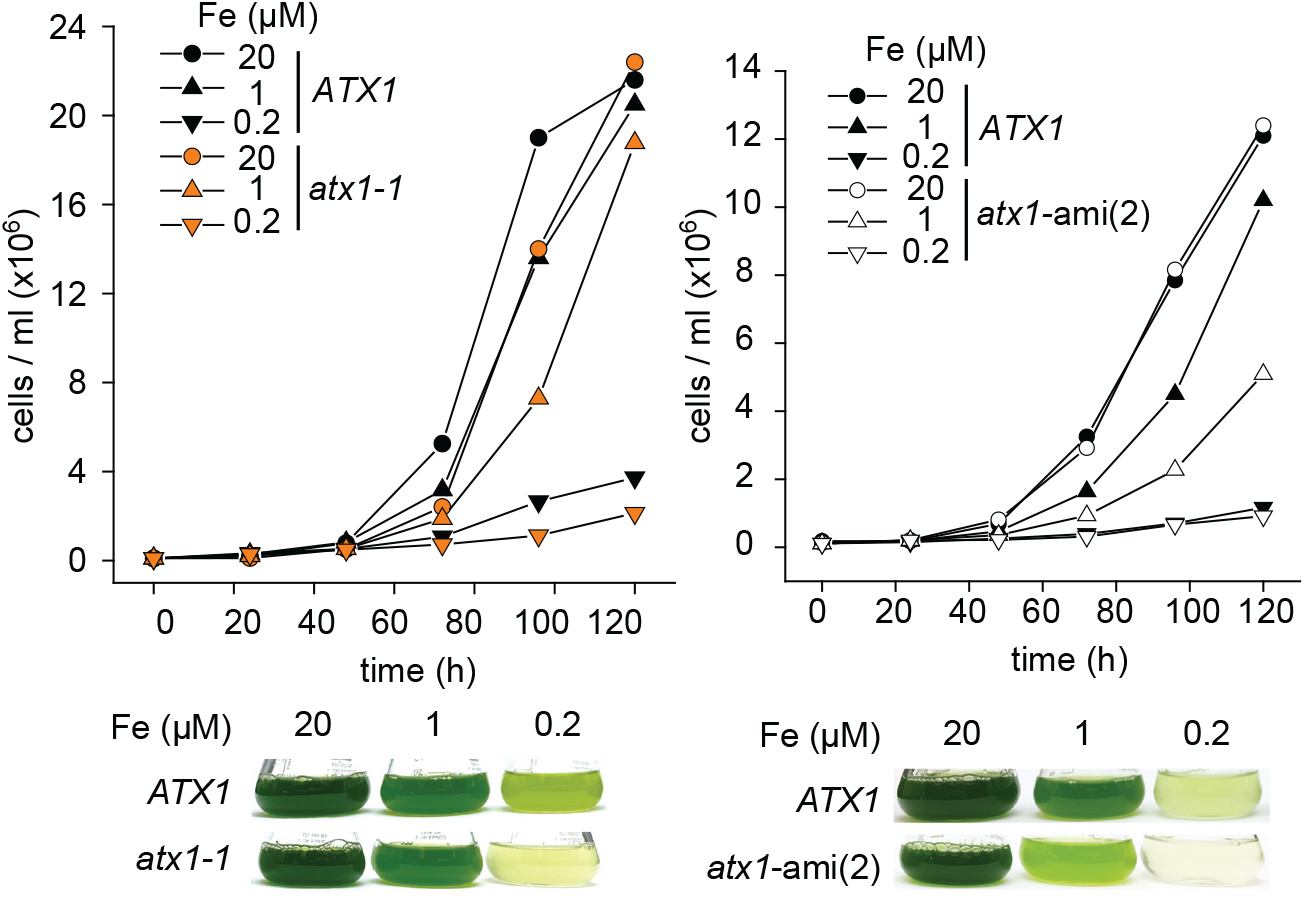
Iron nutrition dependent growth defect is exacerbated in *atx1* mutants and *atx1*-ami lines. *ATX1* reference lines (black, filled symbols), *atx1* mutants (orange filled symbols) and *atx1*-ami(2), white symbols) were grown in either iron replete TAP medium (20), in Fe deficient (1) or in Fe limited (0.2) growth medium as indicated. Cell counts were obtained at 24-hour intervals using a Coulter counter. Below each panel are pictures of flasks taken 6 days post inoculation.

**Figure 7:**
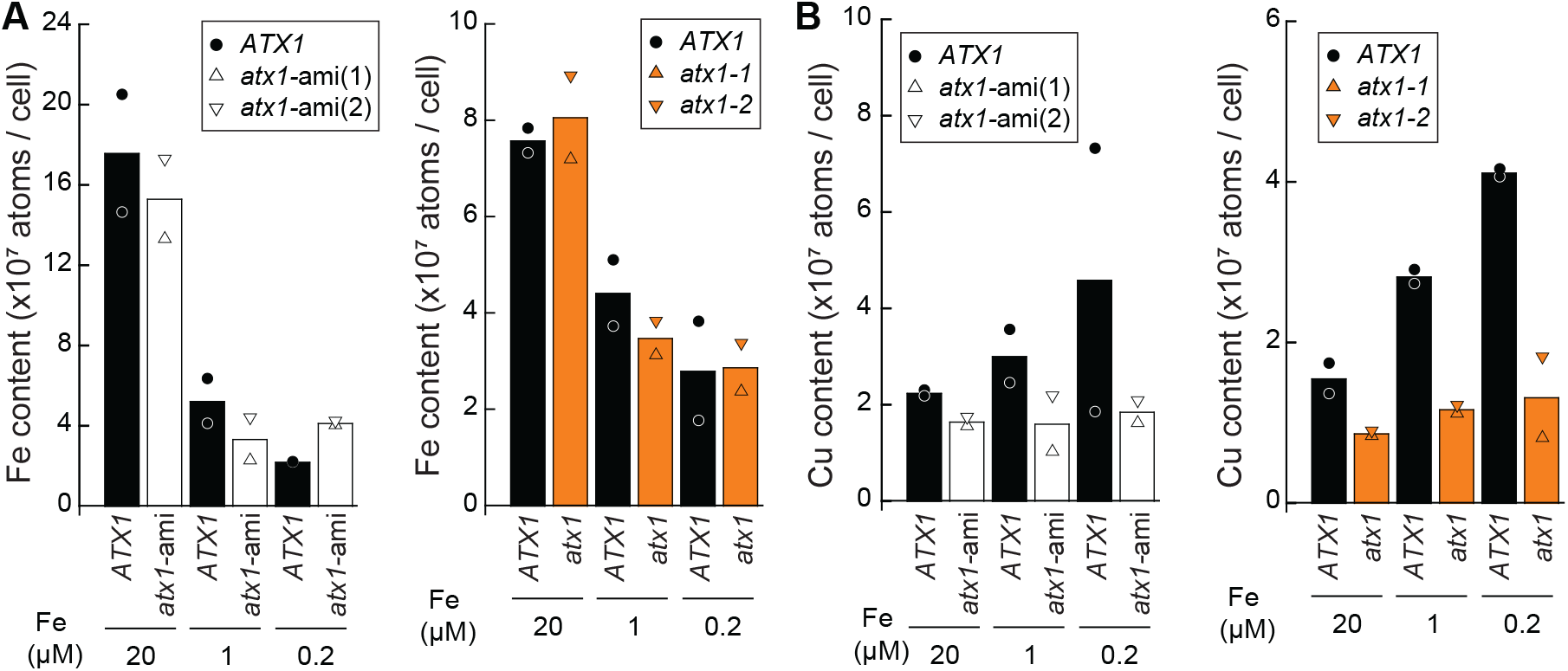
Iron and copper content in *atx1* mutants and *atx1*-amiRNA lines. **(A)** Iron and **(B)** copper content of empty vector (*ATX1*, black) and *atx1*-ami lines (*atx1*-ami(1) and *atx1*-ami(2), white triangles), mutant background strains (ATX1, black), *atx1* mutants (*atx1-1* and *atx1-2*, orange triangles), that were grown in media containing iron as indicated (replete 20, deficient 1, and limited 0.2) was measured by ICP-MS. Shown are averages and individual values from independent grown cultures.

We noticed before that Fe deficient cells slightly up-regulate components of the Cu assimilation pathway, likely due to a link between both pathways via FOX1 (Dancis et al., 1994). Accordingly, the copper content of cells grown on growth medium lacking iron is increased 2-fold compared to iron replete grown cells (Figure 7B). On the other hand, Cu content was significantly lower in ami-*atx1* lines as well as *atx1* mutants as compared to their respective reference lines in all conditions that were analyzed (Figure 7B). These data indicate a second phenotype in *atx1* mutants, namely a remarkable impact on Cu assimilation / accumulation.

### ATX1 is important for purine assimilation in nitrogen poor conditions

Urate oxidase (UOX1), an enzyme involved in purine metabolism, is also a Cu enzyme making it another candidate client of ATX1. In accordance with our expectations, we noted that *UOX1* and *ATX1* transcripts both increase transiently under N-deficiency (Figure 8A). To test whether UOX1 is an ATX1 client, we grew the reference wild-type strains and each *atx1* mutant on guanine vs. allantoin as a sole nitrogen source (Figure 8B). Both Guanine and allantoin are taken up by Chlamydomonas, but guanine metabolism requires UOX1 while allantoin metabolism does not (Lisa et al., 1995; Merchán et al., 2001; Piedras et al., 1998). Strikingly, after 72h of growth, *atx1* mutants showed a visible growth defect on guanine but not allantoin as compared to the corresponding reference lines (Figure 9A), with significant less biomass accumulation as determined by chlorophyll content (Figure 9B). We conclude that ATX1 is required for UOX1 function, most likely for its metalation.

**Figure 8:**
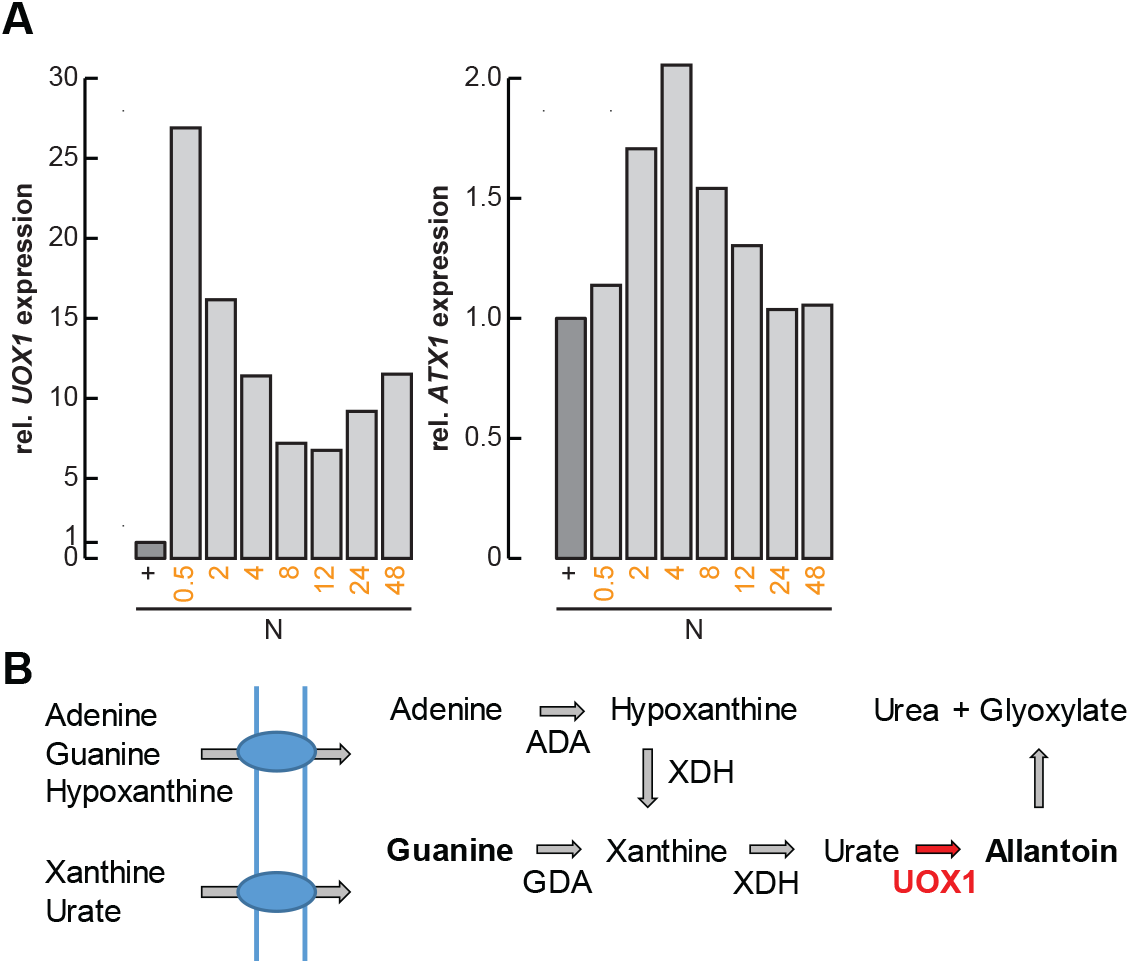
*UOX1* and *ATX1* are transiently expressed upon transfer to N-free medium. **(A)** Relative abundance changes of *UOX1* (left panel) and *ATX1* (right panel) transcripts upon nitrogen limitation. **(B)** Shown is the nitrogen assimilation pathway from purines towards urea. ADA, Adenine deaminase; GDA, Guanine deaminase; XDH, xanthine dehydrogenase; UOX1, urate oxidase; AC, allantoicase

**Figure 9:**
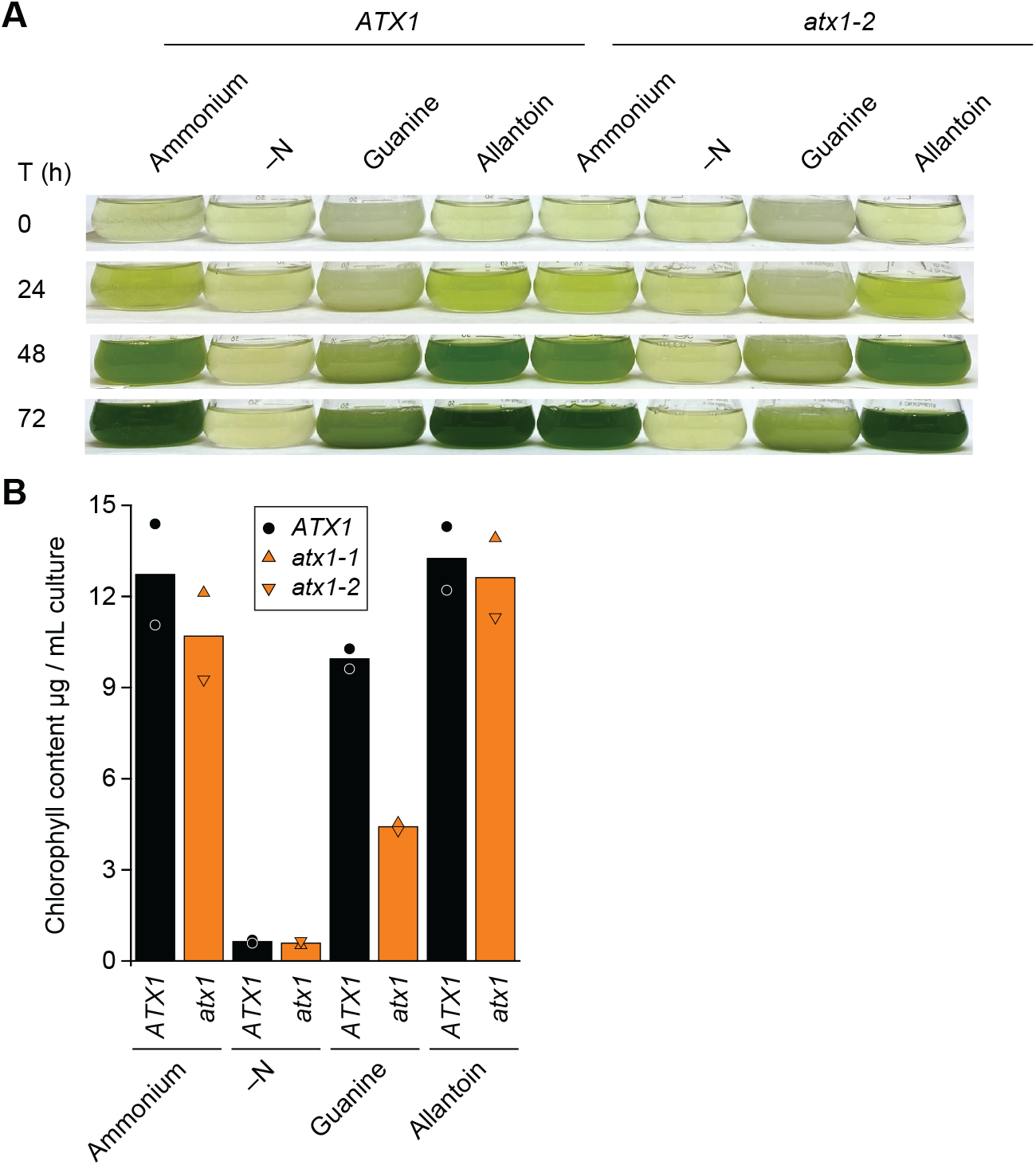
Guanine dependent growth defect in *atx1* mutants. **(A)** Shown is the nitrogen assimilation pathway from purines towards Urea. ADA, Adenine deaminase; GDA, Guanine deaminase; XDH, xanthine dehydrogenase; UOX1, urate oxidase; AC, allantoicase. **(B)** Reference lines and *atx1* mutants were grown in medium lacking any nitrogen source (–N) or supplemented with ammonium, guanine or allantoin as sole nitrogen source as indicated. Pictures of flasks were taken in 24 h intervals. **(C)** Shown is the chlorophyll content of all strains and conditions at time 72 h. Shown are averages and individual values from independent grown cultures as indicated.

## Discussion

### CrATX1: a route for Cu towards the secretory pathway?

In Chlamydomonas, multiple pathways of Cu trafficking emerge from the high affinity, CTR-type Cu import system (Page et al., 2009). Cu(I) taken up by CTRs is delivered to cytosolic Cu chaperones, perhaps via glutathione as an intermediate carrier (Miras et al., 2008), for re-distribution to the mitochondria, the chloroplast and to the secretory pathway (Valentine and Gralla, 1997). The work presented herein describes a soluble Cu chaperone, CrATX1, that we propose to function in Cu delivery to secretory compartments. Several lines of evidence support a role for CrATX1 in Cu sequestration from the inner surface of the plasma membrane region to the secretory pathway, where Cu is subsequently provided to the multicopper Fe (II) oxidase FOX1, the urate oxidase UOX1 and perhaps other secretory Cu containing target proteins awaiting maturation and metalation.

First, both FOX1 and ATX1 are conditionally expressed as a function of iron nutrition. Cells grown in iron limited media are characterized by chlorosis and growth arrest and this phenotype is exacerbated in *atx1* mutants. Yet, strains with reduced or even no ATX1 abundance have the same iron content as do wild type cells. This phenotype mirror that noted in *fox1* knock down lines (Chen et al., 2008). This is because Fe is a growth limiting nutrient. When the minimal Fe quota is not met, no further growth can occur, and ami-*atx1* lines as well as *atx1* mutants recapitulate the iron conditional growth defect of *fox1* knock-down lines. Since both, *fox1* and *atx1* mutant strains do grow, albeit poorly, in iron-poor conditions, we propose that there must be a Cu independent alternative system to facilitate high affinity iron uptake. The ZIP family transporters IRT1 and IRT2 are likely candidates (Blaby-Haas and Merchant, 2012; Chen et al., 2008). This is further supported by the observation that Cu deficient cells are also not secondarily iron deficient, despite FOX1 lacking its Cu co-factor in this situation (Kropat et al., 2015). Second, we see a growth phenotype when cells are grown in media with guanine as the sole nitrogen source in which they are dependent on purine assimilation via UOX1, which is also metalated in the secretory pathway. Again, the *atx1* mutants show reduced growth as compared to reference wild-type lines. The third line of evidence is the reduced copper content of *atx1* mutants, which speaks to the role of the chaperone in Cu(I) assimilation.

In addition, by expressing an N terminal YFP-ATX1 fusion protein in Chlamydomonas, we were able to demonstrate that CrATX1 is localized within the cytoplasm and more specifically seems to be localized to the inner plasma membrane and to the trans-Golgi Network (Figure 4B). These results are consistent with the function of ATX1 in other organisms. ATX1 present within the cytoplasm receives Cu from CTRs and GSH, and based on fluorescence imagery of cells expressing the N terminal YFP-ATX1 fusion protein, ATX1 then concentrates in the trans-Golgi Network via the secretory pathway. ATX1 delivers Cu to target proteins within the trans-Golgi Network, including Cu P-type ATPases such as the ATP7A homologue, CTP1, that likely feeds Cu into the Golgi lumen and whose expression is also induced in iron starved cells. Loading of the multicopper oxidase, FOX1, as well as of UOX1 is dependent on Cu delivered via this pathway.

## Acknowledgments

This work was supported by NIH grant GM42143 (to S.S.M.). Confocal microscopy experiments were conducted using a Zeiss LSM 880 with OPO, at the CRL Molecular Imaging Center, supported by the Helen Wills Neuroscience Institute. We would like to thank Holly Aaron and Feather Ives for their microscopy training and assistance, and Chris Jeans and the QB3 Macrolab at UC Berkeley for purification of LbCPf1.

