## Supplemental Figures 1-4 for "Chlamydomonas ATX1 is essential for Cu distribution towards the secretory pathway and maintenance of biomass in conditions demanding cupro-enzyme dependent metabolic pathways"

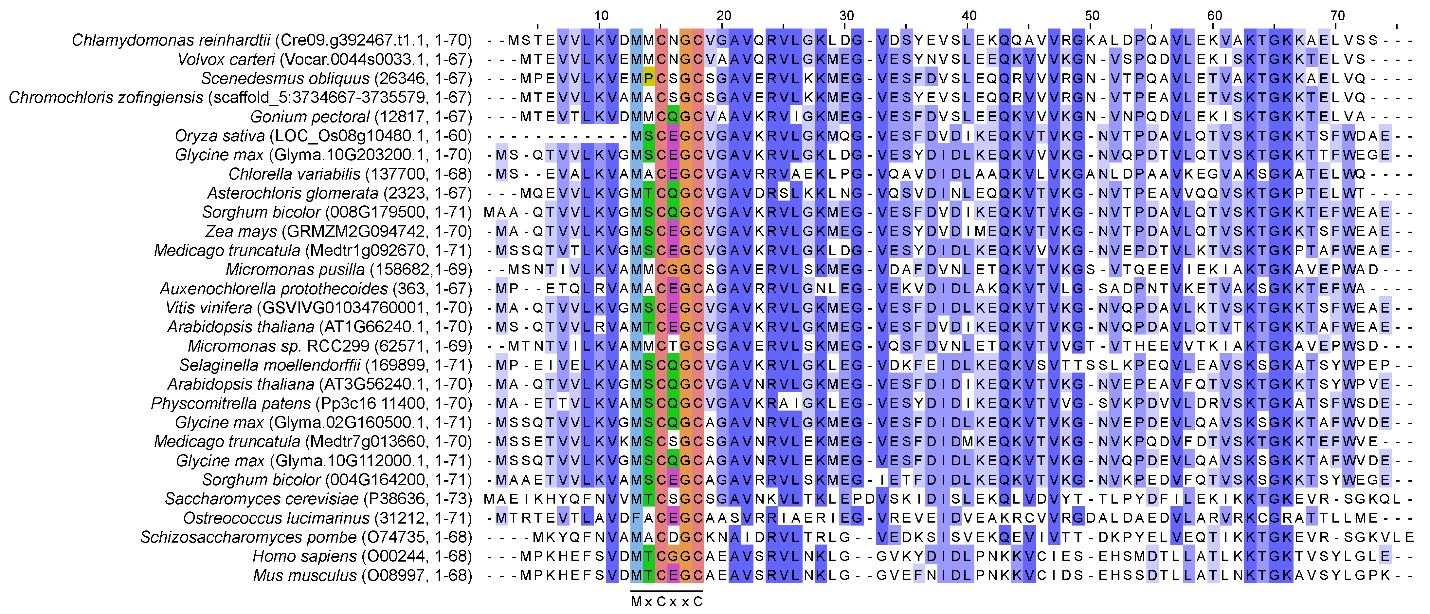
**Supplemental Figure 1**. Shown is a multiple sequence alignment using Atx1 protein sequences from diverse organisms.

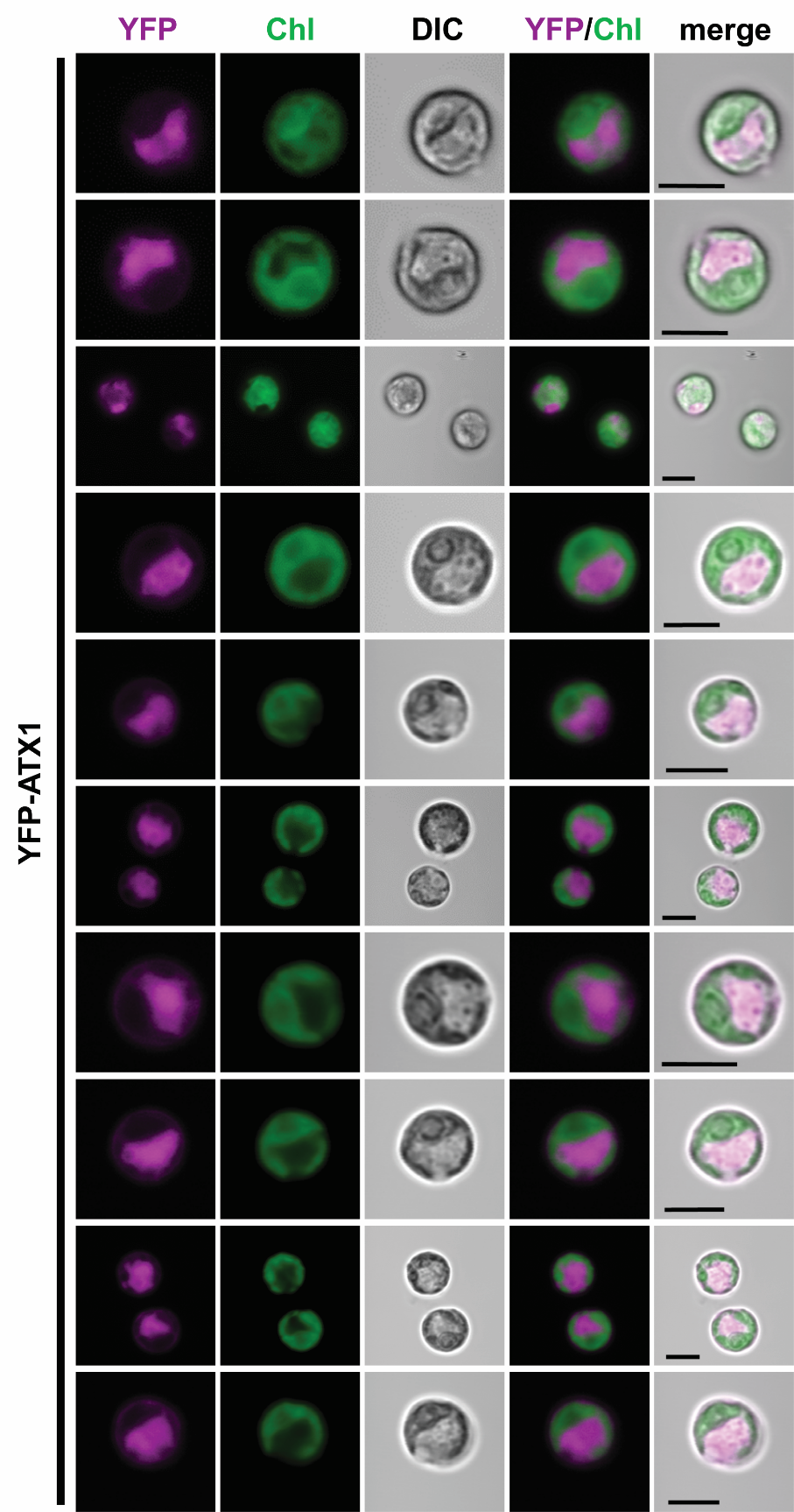

**Supplemental Figure 2.** The YFP-ATX1 fusion protein localizes to the cytosol. Figure shows all cells that were imaged in experiments shown and described in Figure 4A.

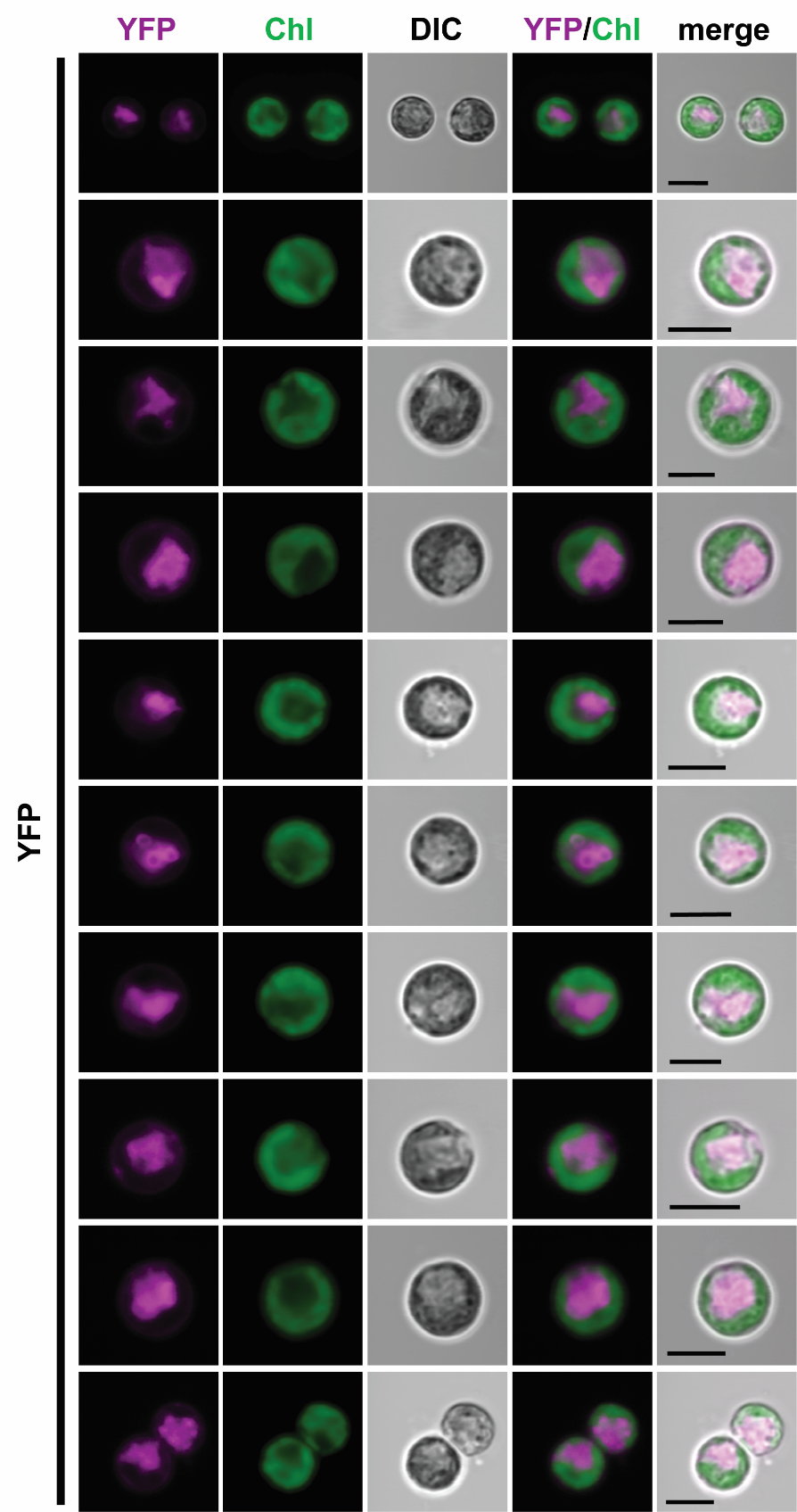

**Supplemental Figure 3**. YFP localizes to the cytosol. Figure shows all cells that were imaged in experiments shown and described in Figure 4A.

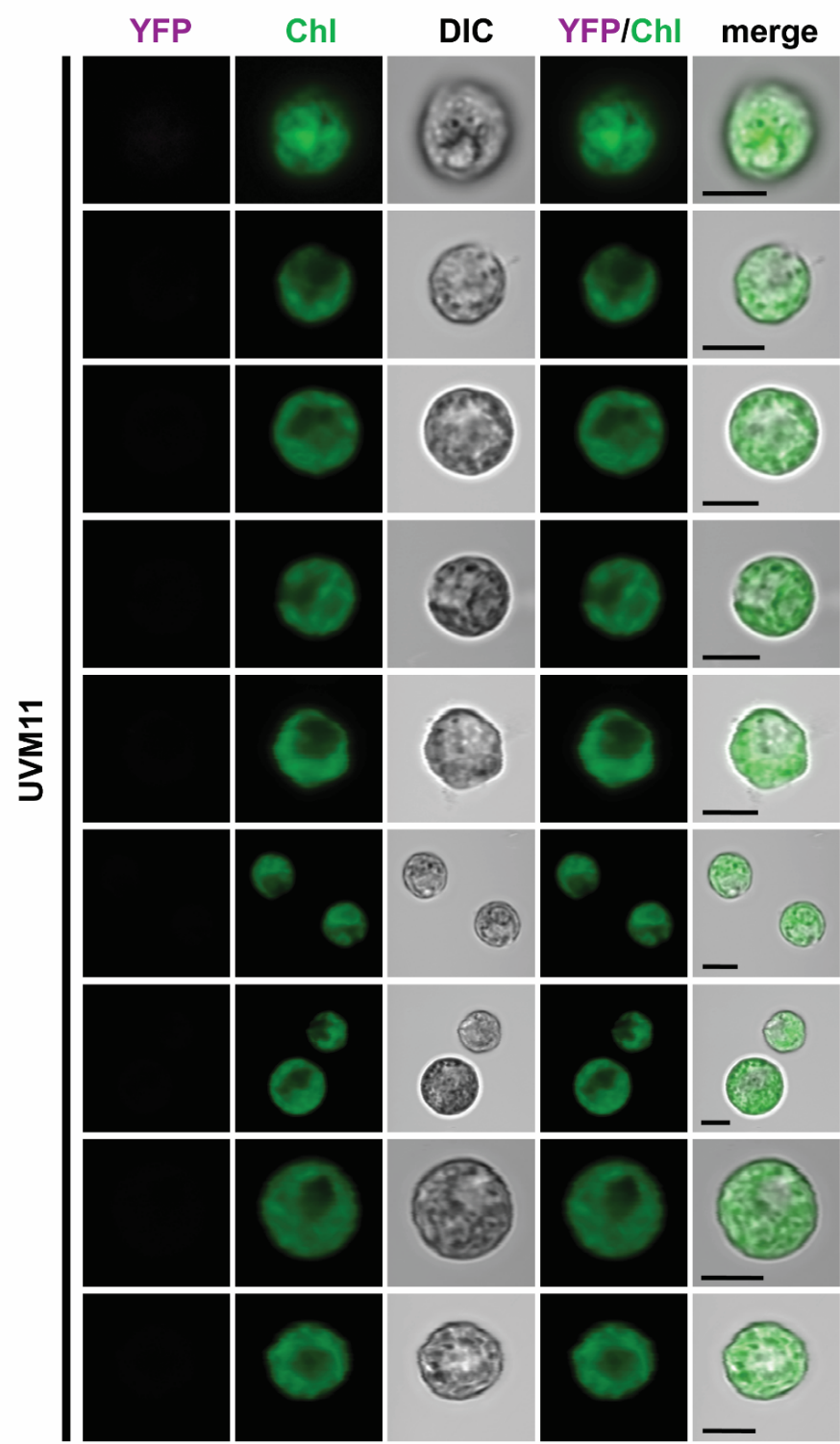

**Supplemental Figure 4.** No YFP signal was detected in the UVM11 background strain. Figure shows all cells that were imaged in experiments shown and described in Figure 4A.
